# Neuronal network activity controls microglial process surveillance in awake mice via norepinephrine signaling

**DOI:** 10.1101/557686

**Authors:** Yong Liu, Yujiao Li, Ukpong B. Eyo, Tingjun Chen, Anthony D. Umpierre, Jia Zhu, Dale B. Bosco, Hailong Dong, Long-Jun Wu

## Abstract

Microglia are resident immune cells that dynamically survey the brain parenchyma, interacting with neurons in both health and disease. However, it is still unclear how neuronal network activity drives microglial dynamics. Utilizing *in vivo* two-photon imaging of microglia in awake mice, we found that inhibition of neuronal activity under general anesthesia dramatically increased microglial process surveillance. Accordingly, both sensory deprivation and optogenetic inhibition of local neuronal network activity in awake mice resulted in similar increases in microglial process surveillance. We further determined that reduced norepinephrine signaling is responsible for the observed increase in microglial process surveillance. Our results demonstrate that microglial process dynamics are directly influenced by neural activities through norepinephrine signaling in awake animals and indicate the importance of awake imaging for studying microglia-neuron interactions.

## Main text

Microglial processes dynamically survey the brain parenchyma, extending, and converging their processes towards sites of brain injury or neuronal hyperactivity (*1-3*). Multiple studies demonstrate that microglial process dynamics play an important role in maintaining brain homeostasis in health and diseases. However, microglial process dynamics *in vivo* have been studied predominantly under anesthesia, and it is unclear whether and how neuronal network activity under awake condition may affect microglia behaviors. Here, we utilized *in vivo* imaging techniques to examine microglia process surveillance in the awake mice and compared those dynamics to periods of acute anesthesia, sensory deprivation, or optogenetic inhibition.

We visualized and quantified microglia process surveillance using CX3CR1^GFP/+^ mice under in vivo two-photon microscopy. After cranial window surgery, mice were trained to freely move on a custom-made rotating treadmill while head restrained (Fig. S1A-C). Under awake conditions, we observe that microglia processes maintain a relatively stable territory of surveillance (Fig. S1D-F). We hypothesized that microglia process surveillance has a partial dependence on neuronal activity, and thus would be attenuated by general anesthesia. When isoflurane (1.2% in oxygen, 700 mL/min) was applied to the animal via nose cone, we surprisingly observed that microglial processes started to extend and survey greater territory with increased velocity, while the neuronal network activity was suppressed illustrated by dampened neuronal calcium spikes (Fig. 1A-D, Fig. S2A-D and movie S1 and S2) and presented with slow wave activity (Fig. S3E and F). The extension and velocity of microglia processes peaked within 20 min after anesthesia initiation. Sholl analysis confirmed the increase of microglia process length and branch points in the anesthetized mouse (Fig. 1E-G). These results are unexpected as isoflurane has been previously reported to reduce microglial basal motility via inhibition of microglial two-pore domain channel THIK-1 (*4*). In addition, neuronal hyperactivity following acute seizure induction trigger microglia process extension and/or convergence (*5*, 6). Therefore, to test whether increased microglia process surveillance is specific to isoflurane anesthesia or generalizable across multiple anesthetic agents, we examined two other commonly used general anesthetics, ketamine/xylazine (87.5 mg/kg Ket, 12.5 mg/kg Xy) and urethane (1.6 g/kg). Similar to isoflurane inhalation, both ketamine/xylazine and urethane increased microglia process dynamics (Fig. 1H and I). Together, these results reveal that general anesthetics promote rather than inhibit microglial surveillance, suggesting microglial surveillance may not be lower than previously thought under the most physiologically relevant conditions.

**Figure 1.**
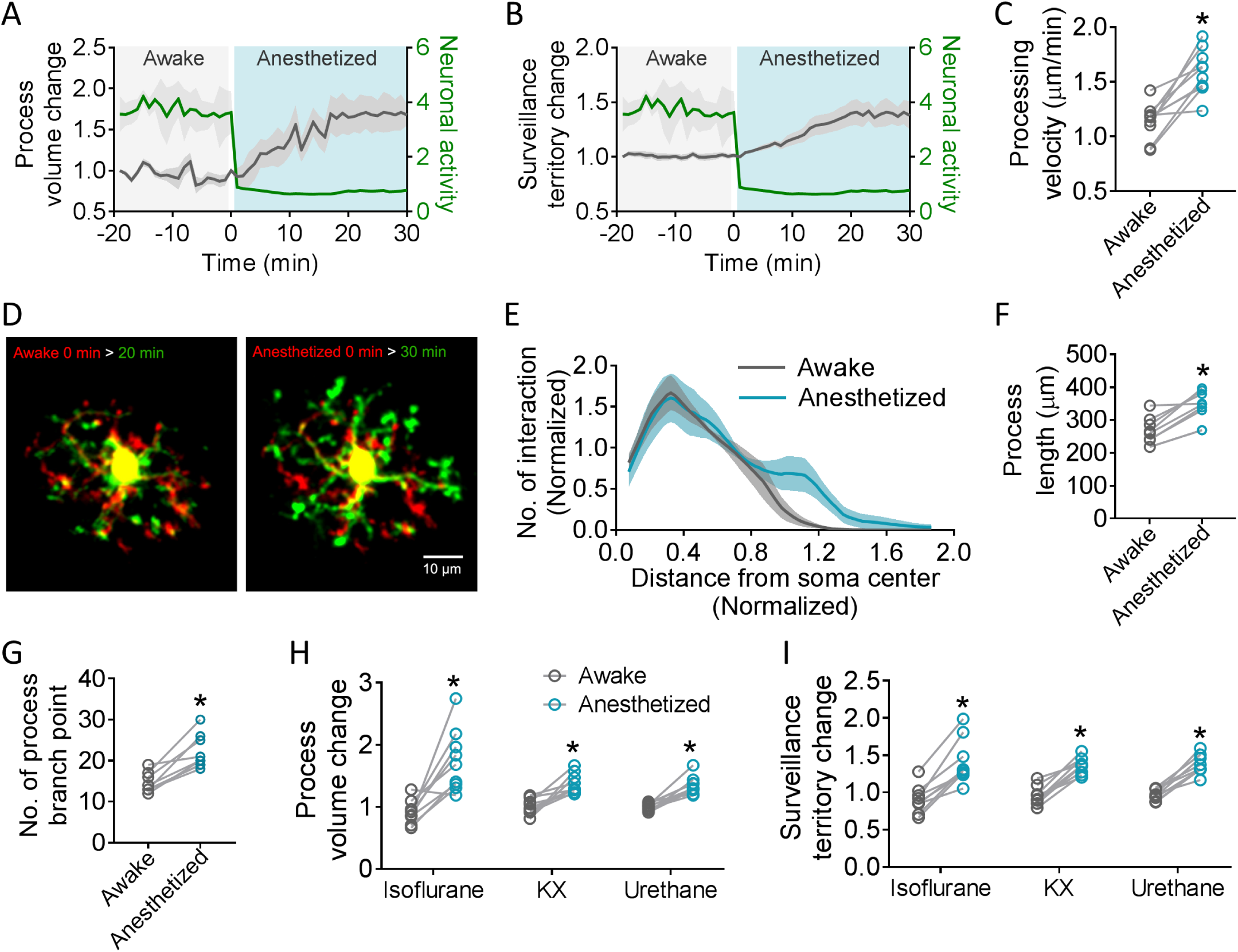
Microglial process dynamics are increased after general anesthesia. (A and B) Time-lapse changes of microglial process volume (A) and surveillance territory (B) and neuronal activity indicated by calcium oscillations (see method for calculation) going from an awake to an anesthetized condition. (C) Process velocity changes under awake and anesthetized conditions. 11 cells from 3 mice studied across the two conditions. (D) Representative images of a microglial cell during awake (left panel) and anesthetized (right panel) conditions. Color overlays show initial process territories (red, 0 min) and their change at the end of a 20 min period (green) under the two conditions. (E) Sholl analysis of microglial morphology before and after anesthesia. (F and G) Length (F) and branch points number (G) of microglial process before and after anesthesia. (A-G) Anesthesia were initiated and maintained by isoflurane. (H and I) Microglial process volume (H) and surveillance territory (I) changes induced by different anesthetic agents: isoflurane, ketamine/xylazine (KX), and urethane in awake mice. All points represent 9 cells from 3 mice each group, \**P*≤0.001, paired t-test.

General anesthesia not only reduces neuronal activity, but also alters metabolism, blood pressure and cardiac output (*7*, 8). Thus, to mitigate these adjacent effects, we employed unilateral whisker trimming as a sensory deprivation approach (*9*). Whisker trimming suppresses neuronal activity in the contralateral barrel cortex as shown by GCaMP6 imaging (Fig. S3, movie S3). We then compared microglial process surveillance before and after whisker trimming. Our results showed that whisker trimming triggered increased microglial process volume and surveillance territory in the barrel cortex similar to that induced by general anesthesia (Fig. 2A and B and movie S4). Furthermore, sensory deprivation only increased process surveillance in contralateral barrel cortex, with no effects observed in the ipsilateral cortex (Fig. 2C and D). Whisker trimming further suggests that microglial process surveillance can be triggered by suppression of neuronal activity within a sensory network. To determine whether decreased neuronal activities under sensory deprivation and general anesthesia employ common or distinct mechanisms in triggering increase of microglial process dynamics, we anesthetized mice with isoflurane 30 min after whisker trimming. Microglial process surveillance did not increase further when whisker trimming was followed by isoflurane inhalation (Fig. 2E), indicating that isoflurane does not have an additive effect. Such a finding suggests that sensory deprivation via whisker trimming and general anesthesia may share similar mechanisms for enhancing microglial process surveillance following neuronal activity suppression.

**Figure 2.**
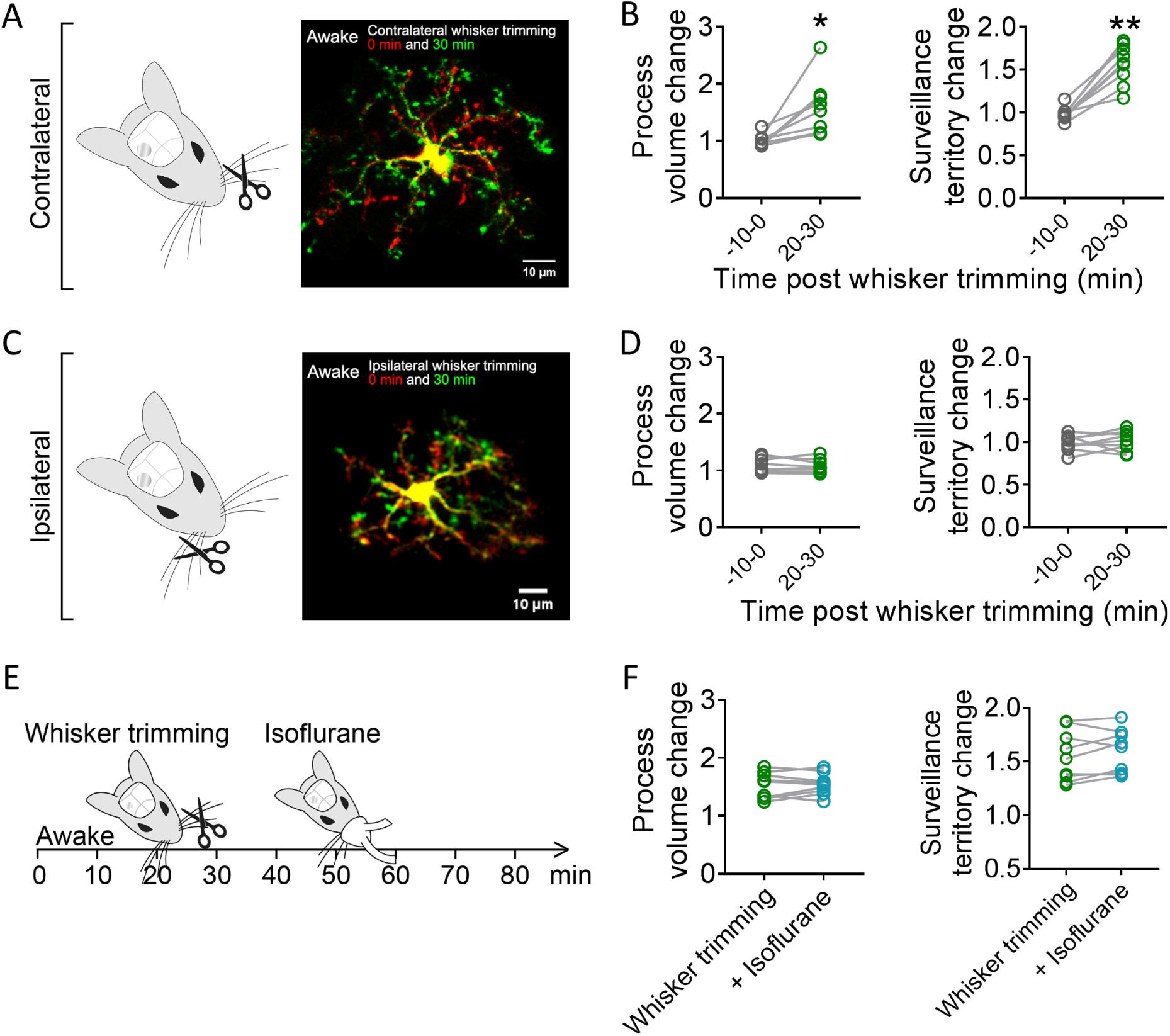
Whisker trimming increases microglial process surveillance in the barrel cortex of awake mice. (A) Whisker trimming and imaging of microglia in the contralateral barrel cortex before (red) and 30 min after (green) whisker trimming in awake mice. (B) Increased microglial process volume and surveillance territory after trimming of contralateral whisker. (C) Whisker trimming and imaging of microglia in the ipsilateral barrel cortex before (red) and 30 min after (green) trimming in awake mice. (D) No observed changes in microglial process dynamics before and after ipsilateral whisker trimming. (E) Experimental paradigm: microglial process dynamics were studied in the awake animal, followed by whisker trimming at 20 minutes later. An additional 30 minutes after whisker trimming, isoflurane anesthesia is applied to the mouse. Microglial process dynamics in the contralateral barrel cortex following trimming and anesthesia are normalized to the awake condition and do not suggest additive effects of anesthesia following whisker trimming. All points represent 9 cells from 3 mice, \**P*<0.01, \*\**P*<0.001, paired t-test.

To directly manipulate neuronal activity and test its effects on microglial process surveillance, we intracerebrally applied muscimol (870 µm), a potent and selective agonist for GABA_A_ receptors (*10*) in awake mice. Muscimol had the ability to strongly inhibit neuronal activity showing by Ca^2+^ imaging in mice (Fig. S4 and movie S5). Similar to those under general anesthesia and sensory deprivation, muscimol application increased microglia process surveillance in awake mice (Fig. 3A and B, movie S6) relative to intracerebral aCSF application (Fig. 3C). Next, we employed an optogenetic strategy to suppress neuronal activity in real time. We expressed channelrhodopsin 2 (ChR2) in VGAT-positive GABAergic neurons (VGAT-ChR2) to specifically inhibit neuronal networks on demand. The expression of ChR2 in VGAT-positive neurons was confirmed by electrophysiology in acute brain slices (Fig. 3D and E and S5). In awake mice, we applied LED light pulses (470nm, 10 Hz, 10 min) to optogenetically activate GABAergic neurons and then compared microglial process surveillance before and after optogenetic inhibition of the somatosensory cortex. Following GABAergic stimulation, we indeed observed an increase in microglial process surveillance in awake mice (Fig. 3F-H and movie S7). Together, these results demonstrate that inhibition of neuronal network activity by pharmacological or optogenetic approaches increases microglial process surveillance in awake mice.

**Figure 3.**
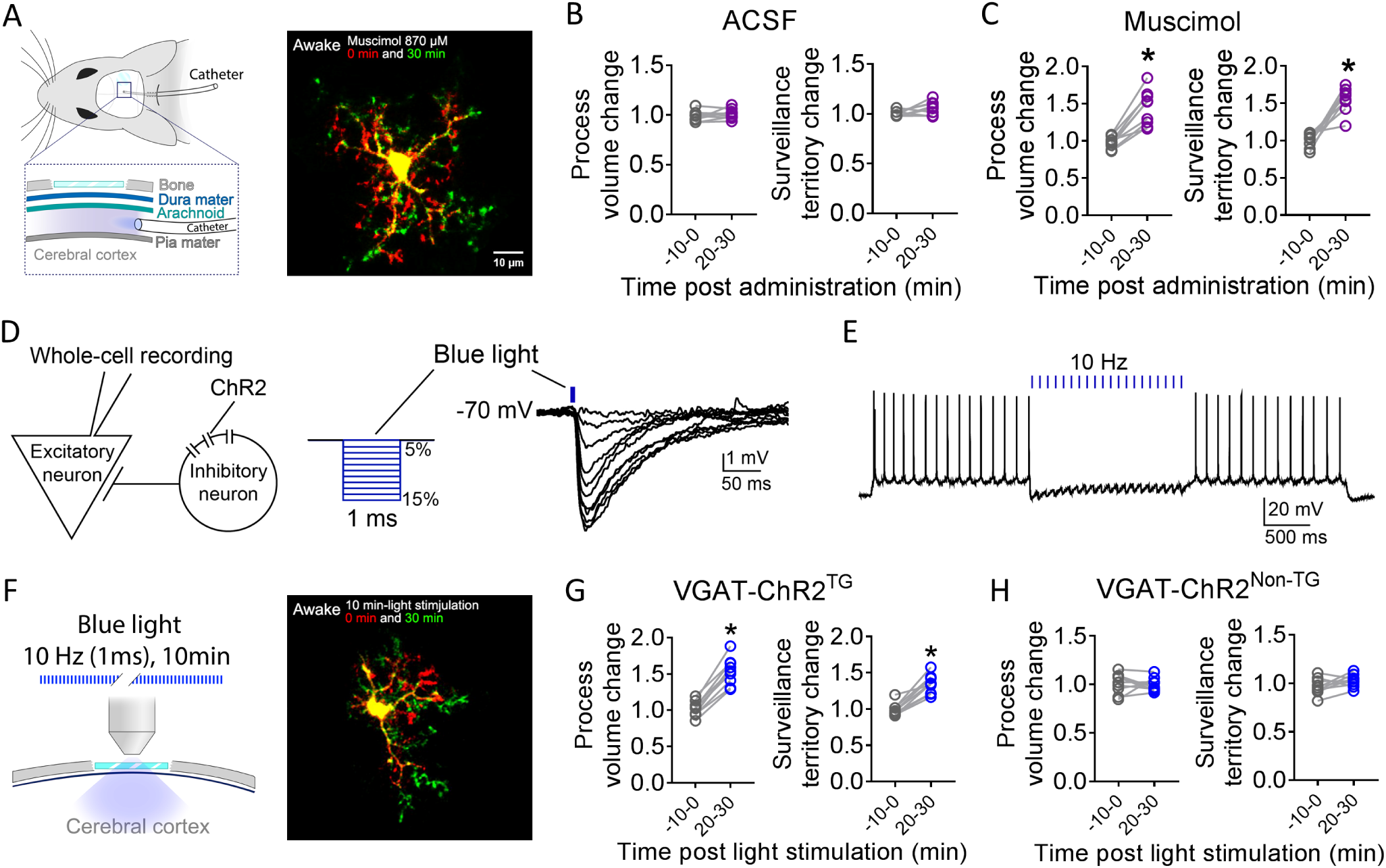
Suppression of neuronal network activity by pharmacological and optogenetic approaches increases microglial process surveillance. (A) Imaging of microglia before (red) and 30 min after (green) intracerebral administration of muscimol. (B and C) Changes in microglial process volume and surveillance territory before and after administration of ACSF (B) or muscimol (C). (D) Whole cell recording of excitatory neuron identified by firing pattern and morphology in VGAT-ChR2 brain slices (ChR2 expressed in GABAergic neurons). A 1 ms light pulse (470 nm) induced intensity-dependent (5%-15%) membrane hyperpolarization in patched neuron. (E) A 10 Hz pulse of light stimulation inhibited firing of excitatory neurons. (F) Experimental paradigm and imaging of microglia prior to (red) and after (green) 10 min light stimulation. (G and H) Changes in microglial process dynamics before and after light stimulation in VGAT-ChR2^TG^ (G) or VGAT-ChR2^Non-TG^ mice (H). All point represent 9 cells from 3 mice each group, \**P*<0.001, paired t-test.

Next, we wanted to study the molecular mechanism underlying the regulation of microglial process surveillance following neuronal suppression. Because ATP/ADP induces microglial process extension acutely through the P2Y_12_ receptor (*2*, 6), we used P2Y_12_ deficient (P2Y_12_^-/-^) mice to determine if purinergic signaling drives microglial process surveillance when neuronal activities are inhibited. To this end, we performed *in vivo* imaging of microglia in CX3CR1^GFP/+^:P2Y12^-/-^ mice before and after isoflurane anesthesia. Interestingly, increased microglial process surveillance after anesthesia were preserved in P2Y_12_^-/-^ mice (Fig. S6), suggesting that ATP/ P2Y_12_ signaling does not regulate our observed microglia process surveillance under general anesthesia.

Previous studies have found that primary neurotransmitters such as glutamate and GABA cannot directly induce microglial process chemotaxis (*6*, 11), thus we surmised that neuromodulators impacted local neuronal activity (*12*, 13) and microglial dynamics. Acetylcholine, dopamine, serotonin, and norepinephrine (NE) were specifically studied in the context of microglial process dynamics, because the activity of these neuromodulators has been shown to be significantly decreased by general anesthesia (*14*). Interestingly, we found that NE application, but not acetylcholine, dopamine, or serotonin, was able to prevent the increased microglial process surveillance induced by general anesthesia (Fig. 4A-B). These results suggest that reduced NE signaling in anesthetized condition could trigger an increase in microglia process surveillance.

**Figure 4.**
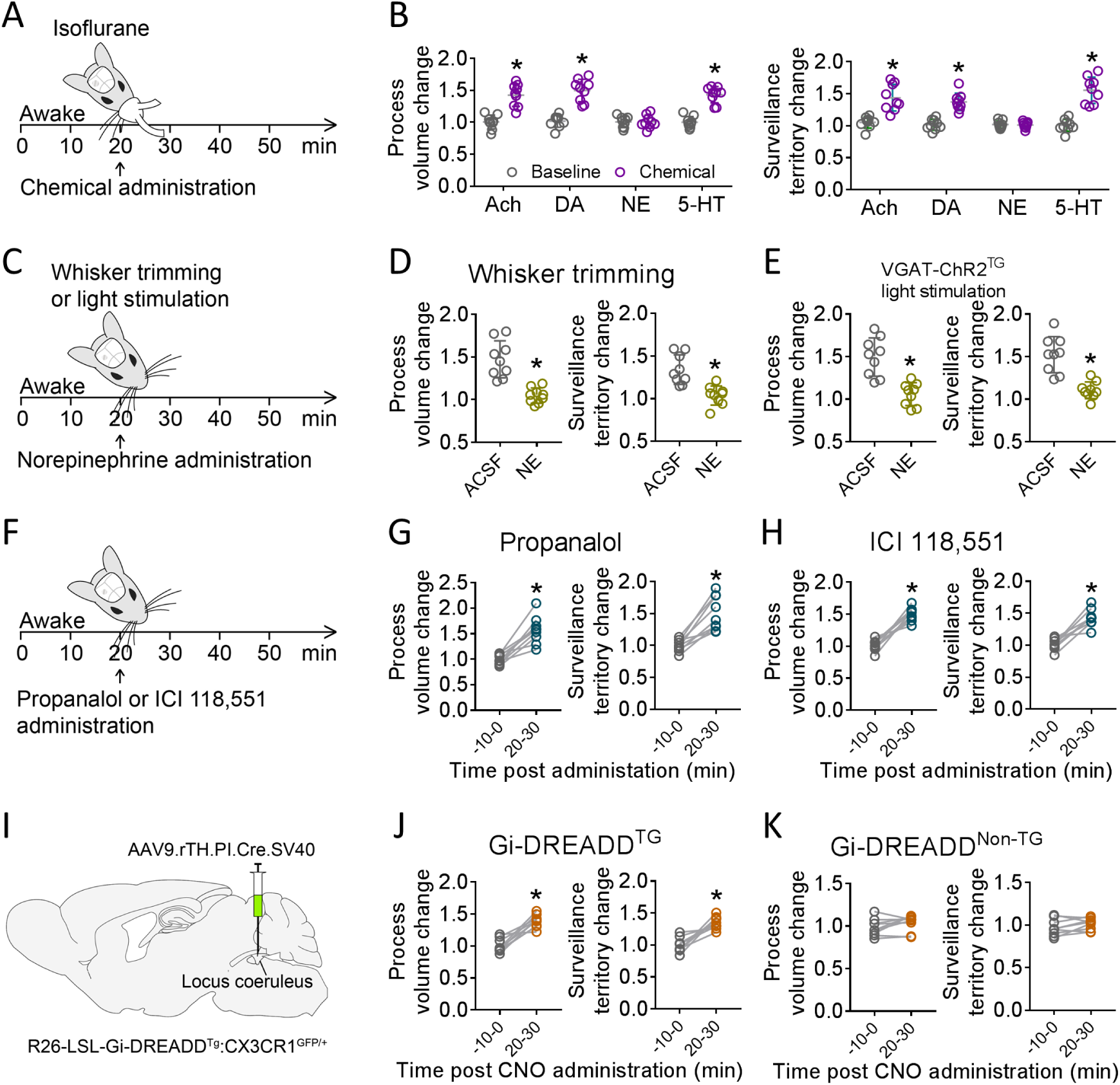
Reduced norepinephrine signaling increases microglial process surveillance after lowering neuronal network activity. (A) Experimental paradigm in (B). (B) Changes of microglial process volume and surveillance territory (microglial process dynamics) in response to isoflurane anesthesia and intracerebral administration of acetylcholine (Ach), dopamine (DA), norepinephrine (NE), or serotonin (5-HT). (C) Experimental paradigm in (D) and (E). (C and D) NE reversed the changes of microglial process dynamics by whisker trimming (C) or by optogenetic inhibition in awake VGAT-ChR2^TG^ mice (E). (F) Experimental paradigm in (G) and (H). (G and H) β-adrenergic receptors blocker propanalol (G) and β2-adrenergic receptors blocker ICI 118,551 (H) increased microglial process dynamics in awake mice. (I-K) Gi-DREADDs were expressed in locus coeruleus adrenergic neurons by indicated virus in the floxed mice (I). CNO increases microglial process dynamics in awake mice expressing Gi-DREADD specifically in locus coeruleus adrenergic neurons (J) but not in control mice without Gi-DREADD expression (K). All points represent 9 cells from 3 mice each group, \**P*<0.001, paired t-test.

We further studied the regulation of microglial process surveillance by NE during whisker trimming or light stimulation in VGAT-ChR2 positive mice (Fig. 4C). Indeed, we found that intracerebral NE administration prevented the increase of microglial process surveillance induced by whisker trimming or optogenetic inhibition (Fig. 4D and E). Microglia are known to express β2-adrenergic receptor that can induce microglial process retraction (*15*). To explore adrenergic receptors that control microglial activity, we intracerebrally applied the β-adrenergic receptor antagonist propanalol or the β2-adrenergic receptor antagonist ICI 118,551 and imaged microglial process surveillance in awake mice (Fig. 4F). We found that either propanalol or ICI 118,551 induced an increase in microglial process dynamics in the awake mouse (Fig. 4G and H). Cortical NE tone is largely generated by adrenergic neurons in the locus coeruleus (LC) (*16*) which project into somatosensory cortex (Fig. S7A and B and movie S8). To directly study the effect of NE signaling on microglia, we used a chemogenetic approach to selectively suppress adrenergic neurons in the LC. An inhibitory Gi DREADD mouse (R26-LSL-Gi-DREADD) was bred with the CX3CR1^GFP/+^ line. Gi-DREADD expression was confined to the LC through targeted AAV-Cre injection (AAV9.rTH.PI.Cre.SV40) (Fig. 4I). We confirmed that the DREADD ligand, clozapine-*N*-oxide (CNO), reduced the firing rate of Gi-DREADD expressing adrenergic neurons in LC (Fig. S7C-F). In the awake mouse, systemic application of CNO (i.p. 5 mg/kg) increased microglia process surveillance in the somatosensory cortex (Fig. 4J and K). Therefore, both chemogenetic inhibition of NE release and β2-adrenergic receptor antagonism increase microglial process surveillance, similar to the effects of general anesthesia, sensory deprivation, or optogenetic inhibition of cortex neuronal activities. Taken together, all categories of our results demonstrated that NE signaling drives microglial process dynamics during reduced neuronal network activity under general anesthesia, sensory deprivation.

Microglia are an immune guardian and shape brain circuit and activity in both health and diseased conditions (*17*). Here, our results revealed that neuronal activity negatively regulates microglial surveillance in awake mice (Fig. S8). When neuronal activity is reduced by various manipulations, such as general anesthesia and sensory deprivation, microglial processes increase dynamics to survey more territory of the brain parenchyma. Supporting this novel observation, a recent study reported that monocular deprivation induced ramification of microglial processes in the contralateral visual cortex (*18*). The current study further demonstrated that enhanced process surveillance following strong reductions in neuronal activity is due to the loss of NE tone and β2-adrenergic receptor activation in microglia. Interestingly, previous research has shown that microglia increase their process dynamics through extension and convergence following periods of neuronal hyperactivity during seizure and stroke contexts (*6*, 19). This increase of microglial process dynamics induced by neuronal hyperactivities is dependent on purinergic signaling and microglial P2Y12 receptors. Therefore, these results suggest that microglia processes are maximally quiescent in the awake animal. Shifting neuronal activity away from this state in either direction can increase process surveillance via different signaling mechanisms. This “set point” pattern of regulation suggests microglial processes are highly attuned to the local environment and respond dynamically to purinergic and noradrenergic signaling.

The increase of microglial process surveillance may suggest the more interactions of microglia with neuronal elements, such axons and dendritic spines (*20*, 21). The function significance of microglial process interaction with neurons is largely unknown. Previous studies indicate the microglial interaction may dampen neuronal activities (*6*, 22), while other studies show that microglial processes increase dendritic spine Ca^2+^ activity (*23*), promote new spine formation during brain development or learning (*21, 24*). In addition, reduced neuronal activity and NE tone also happen under sleep (*25*) or in neurodegenerative disorders (*26*). Therefore, neuronal activity-dependent NE signaling drives the surveying state of microglia processes may contribute to synaptic remodeling in sleep-wake cycle and in neurodegeneration (*27*). In sum, here we provide novel insights of microglia behaviors under awake conditions and their regulation by neuronal activities and NE tone at basal state.

## Acknowledgements

This work is supported by National Institute of Health (R01NS088627, R21DE025689). We thank Dr. Mark Mattson (National Institute on Aging) for critical reading of the paper and members of Wu lab at Mayo for insightful discussions.

## Supplementary Materials

### Materials and Methods

#### Animal

Male and female mice, 2 to 3 months of age, were used in accordance with institutional guidelines. All experiments were approved by the Mayo Clinic Animal Care and Use Committee. Heterozygous (CX3CR1^GFP/+^) mice expressing GFP under the control of the fractalkine receptor (CX3CR1) were used for all experiments (*1*). P2Y_12_ ^-/-^ mice were originally donated by Dr. Michael Dailey at the University of Iowa (Iowa City, IA, USA). Heterozygous R26-LSL-Gi-DREADD (LSL-Gi^TG^, Stock #026219), hemizygous VGAT-ChR2 (Stock #014548) mice were purchased from the Jackson Laboratory (ME, USA). P2Y_12_^-/-^ and R26-LSL-Gi-DREADD (LSL-Gi^TG^) were cross breed with CX3CR1^GFP/+^ for imaging experiments.

#### Virus Injection

For imaging of neuronal network activity *in vivo*, 0.3 ul of AAV9.CamKII.GCaMP6s.WPRE.SV40 virus (#G12289, Penn Vector Core, PA, USA) was injected into the area of somatosensory cortex (2 mm posterior and 1.5 mm lateral to bregma) or barrel cortex (1.45 mm posterior and 3 mm lateral to bregma). To decrease noradrenergic neuronal activity, 0.3 ul of AAV9.rTH.PI.Cre.SV40 virus (#G12845, Penn Vector Core, PA, USA) was injected into the locus coeruleus (5.4 mm posterior, 0.9 mm lateral and 3.7 mm ventral to bregma) of LSL-Gi^TG^: CX3CR1^GFP/+^ mice to initiate the expression of Gi-DREADD.

#### Chronic Window Implantation

Mice were implanted with a chronic cranial window as we previously described(*2*). Briefly, mice were anesthetized with isoflurane (3% induction, 1.2% maintenance) and maintained on a heating pad during surgery. A dental drill (Osada Model EXL-M40) and drill bit (Fine Science Tools, 19008-07) were used to drill open a circular >3 mm diameter window. For the limb/trunk region of the somatosensory cortex, the skull was removed with the center at about 2 mm posterior and 1.5 mm lateral to bregma, while for the barrel cortex, the skull was removed with the center at about −1.45 posterior and ±3 lateral to bregma. A 3 mm glass coverslip previously sterilized in 70% ethanol was placed inside the window, then light curing dental cement (Tetric EvoFlow) was applied around the glass coverslip and cured with a Kerr Demi Ultra LED Curing Light (Dental Health Products). The skull, excluding the region with the window, was then covered with IBond Total Etch glue (Heraeus) and cured with a curing light. Finally, a custom-made head plate was glued with another application of the light curing dental cement to permanently attach the head plate. Mice were allowed to recover from anesthesia on a heating pad for 10 min before they were returned to their home cage. Ibuprofen in drinking water was provided as an analgesic for 48 hours post-surgery. Mice that showed a loss in imaging window clarity before the 2- to 4-week period of observation were not used for this study.

#### *In Vivo* Two-Photon Imaging

All mice with clear chronic glass window were imaged using a two-photon imaging system (Scientifica, UK) equipped with a Ti:Sapphire Mai Tai DeepSee laser (Spectra Physics, CA, USA). Laser wavelength was tuned to 900 nm to image CX3CR1^GFP/+^ microglia, 920 nm to image GCaMP6s fluorescence, or 950 nm to image tdTomato. Imaging utilized a 40× water-immersion lens and a 180×180 μm field of view (512×512 pixels). A 565-nm dichroic mirror with 525-/50-nm (green channel) and 620-/60-nm (red channel) emission filters. The laser power was maintained at 30–40 mW. Imaging in cortex was conducted 50 to 150 μm beneath the brain surface. To image microglial dynamics and neuronal calcium activity in the awake condition, mice were trained to run on a custom-designed circular treadmill (freely rotating disc) while head fixed (Fig S1A). All mice received 3 days of training (2 times a day, 10 min each time) to become familiar with the system prior to imaging. To induce general anesthesia in awake mice, isoflurane (3% for initiation, 1.2% for maintenance) was administrated by an anesthesia nose cone, while ketamine/xylazine (87.5 mg/kg Ket, 12.5 mg/kg Xy) or urethane (1.6 mg/kg) were administered through intraperitoneal injection. To image microglial dynamics, we acquired eight z-stack slices (2 μm step, 14 μm total thickness) every minute (Fig. S1C). For calcium imaging, continuous acquisition (1 Hz) was carried out approximately 60 μm from the brain surface, corresponding to the approximate depth of microglial z-stacks, or at 150 μm depth to study the somatic activity of neurons in cortical layer 2/3.

#### Drug Application

Intracerebral administration of neuromodulators or antagonists was carried out through a catheter implanted 2 weeks after the chronic window implantation surgery (Fig. 3A, left panel). The catheter was filled with ACSF and no microglia activation was observed within the area of the window after implantation. A 10 µL total solution volume was applied through the catheter and into the brain. Neuromodulators were applied at the following concentrations: 5 mM acetylcholine chloride, 1 mM dopamine hydrochloride, 3 mM norepinephrine, 5 mM serotonin hydrochloride. The dose of muscimol chosen (870 µM) represents the minimal dose needed to induce significant suppression of neuronal activity. Clozapine N-oxide (CNO) was intraperitoneally injected at 5 mg/kg to activate Gi-DREADD in awake mice.

#### Data Analysis

Images were analyzed using ImageJ (*3*). For evaluating changes in microglial volume and process surveillance territory under awake and anesthetized conditions, a Z stack was assembled using an image at each Z level with the least distortion in the X-Y plane (selected from five repeat images at each slice level; Fig. S1C). Images in the Z stack were aligned through the StackReg plugin and a maximum-intensity projection (MIP) along the Z axis was created. The MIP image was then thresholded to obtain a two-dimensional rendering of total process area which indicates the microglial process volume. A bounding box that connects the distal-most edges of microglial processes was then created in the X-Y plane. The area encompassed within this bounding box was defined as the microglial surveillance territory. Volume and territory values in subsequent stacks were normalized to the values of the first stack to determine changes over time. Process velocity was measured using the “Manual tracking” plugin. Z stacks acquired 10 to 20 min into imaging were used to evaluate microglial dynamics in the awake condition, while Z stacks acquired 20 to 30 min after anesthesia were used to evaluate microglial dynamics in the anesthetized conditions.

#### Statistical analyses

Statistical analyses were Student’s t test using GraphPad Prism (GraphPad Software, CA, USA). Student Newman-Keuls post hoc tests were applied to detect statistical differences between groups. Data are expressed as mean ± SEM, and a *P* value <0.05 was considered statistically significant.

#### Supplementary Text

**Fig. S1.**
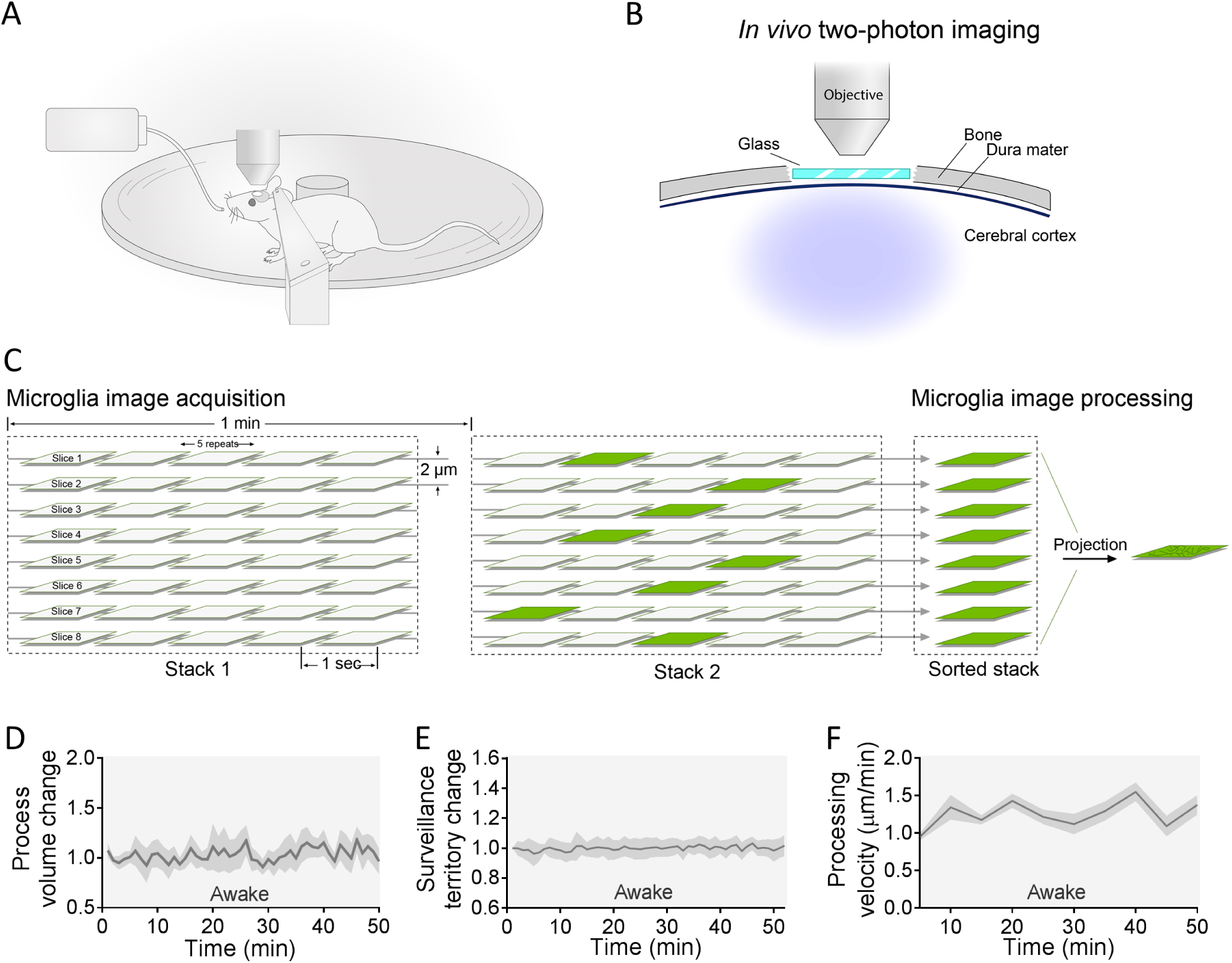
*In vivo* imaging of microglia in awake mice. (A) Head restraint system for mice. (B) Imaging of chronic cranial windows. (C) Microglial image acquisition and processing method for awake mice. (E and E) Time lapse of microglial process volume and surveillance territory in awake mice during a 50 min *in vivo* imaging period. (F) Averaged velocity of microglial processes every 5 min. N= 9 cells from 3 mice/group.

**Fig. S2.**
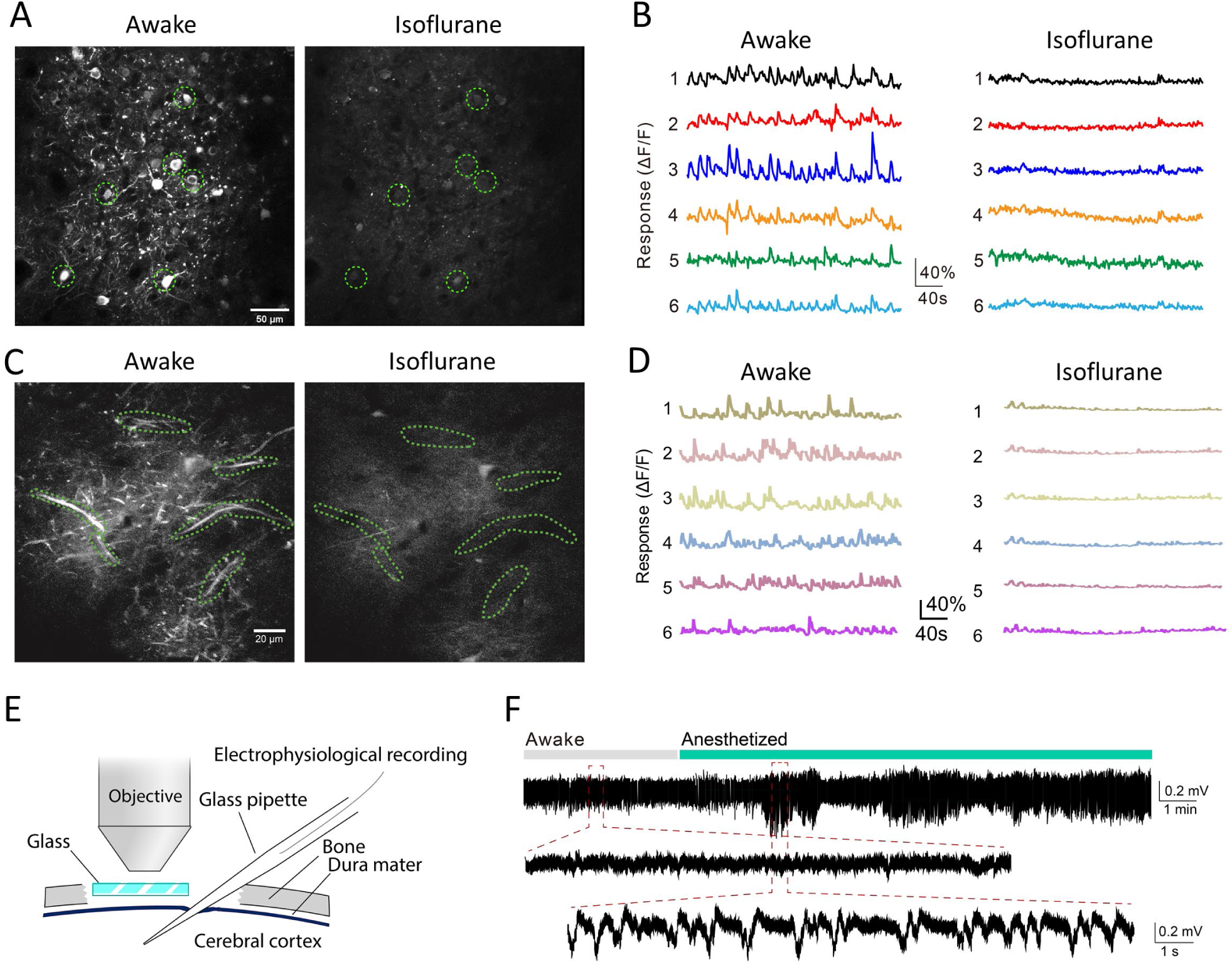
Neuronal network activity in the somatosensory cortex of awake and anesthetized mice. (A-D) Two-photon imaging of neuronal calcium activity in the soma (A) and dendrites (C) of excitatory neurons (soma:150 µm beneath cortical surface; dendrites: 50 µm beneath cortical surface; AAV9.CamKII.GCaMP6s.WPRE.SV40) before and 1 min after isoflurane anesthesia (3% for initiation; 1.2% for maintenance). (B) ΔF/F traces of the six somas circled in (A) before and 1 min after isoflurane anesthesia. (D) ΔF/F traces of six neuronal dendrites highlighted in (C) before and 1 min after anesthesia. (E and F) Field potential recordings 50 µm beneath the cortical surface showing slow wave activity emerging after isoflurane anesthesia.

**Fig. S3.**
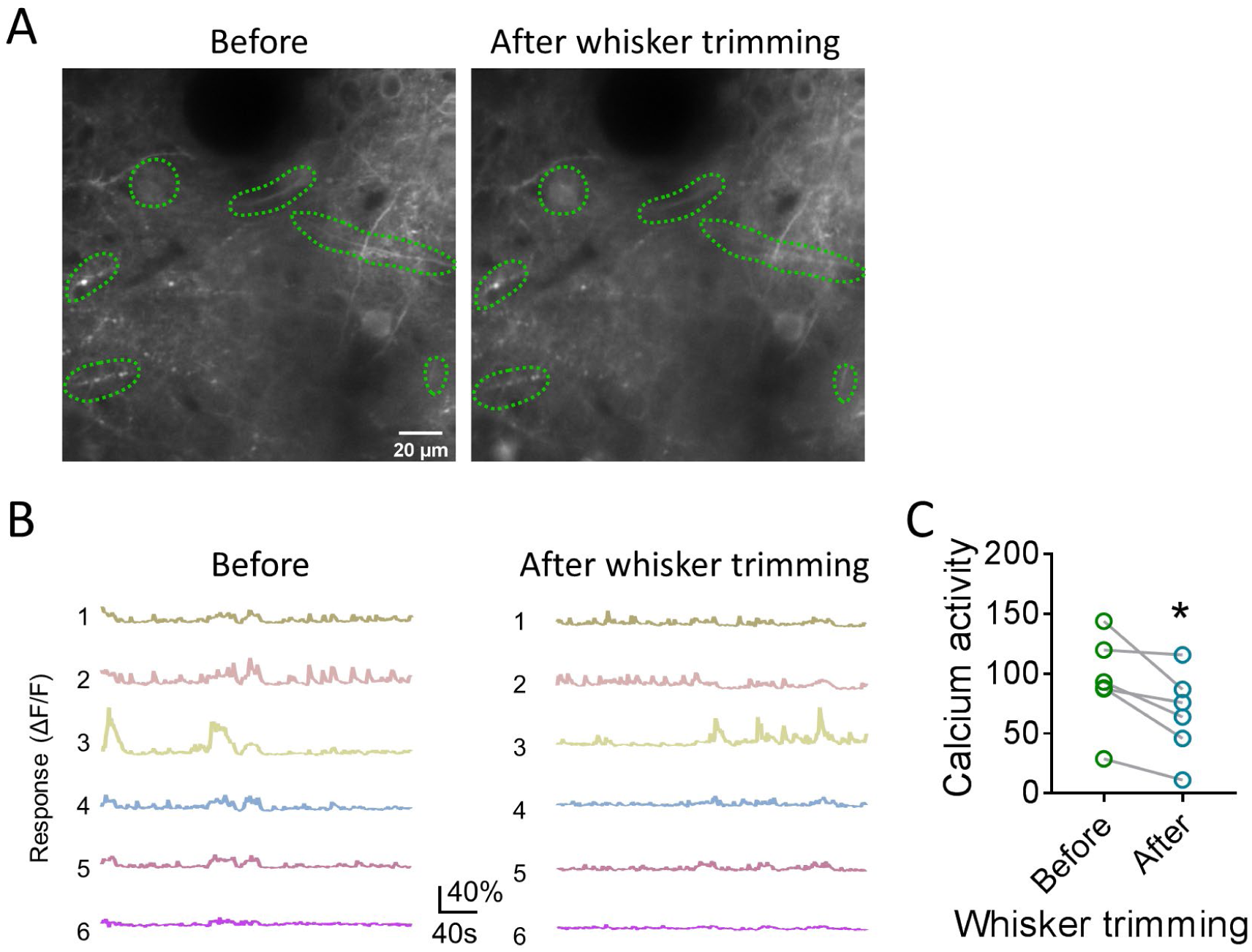
Neuronal network activity decreases after whisker trimming. (A) Two-photon time-lapse images of calcium activity (5 min projection) at 50 µm beneath cortical surface. (B) ΔF/F traces of the six neuronal segments circled in A before and after whisker trimming. (C) Sum of neuronal ΔF/F (value>1.2) in 5 min (calcium activity) before and after whisker trimming. N=6 segments, \**P*<0.05, paired t-test.

**Fig. S4.**
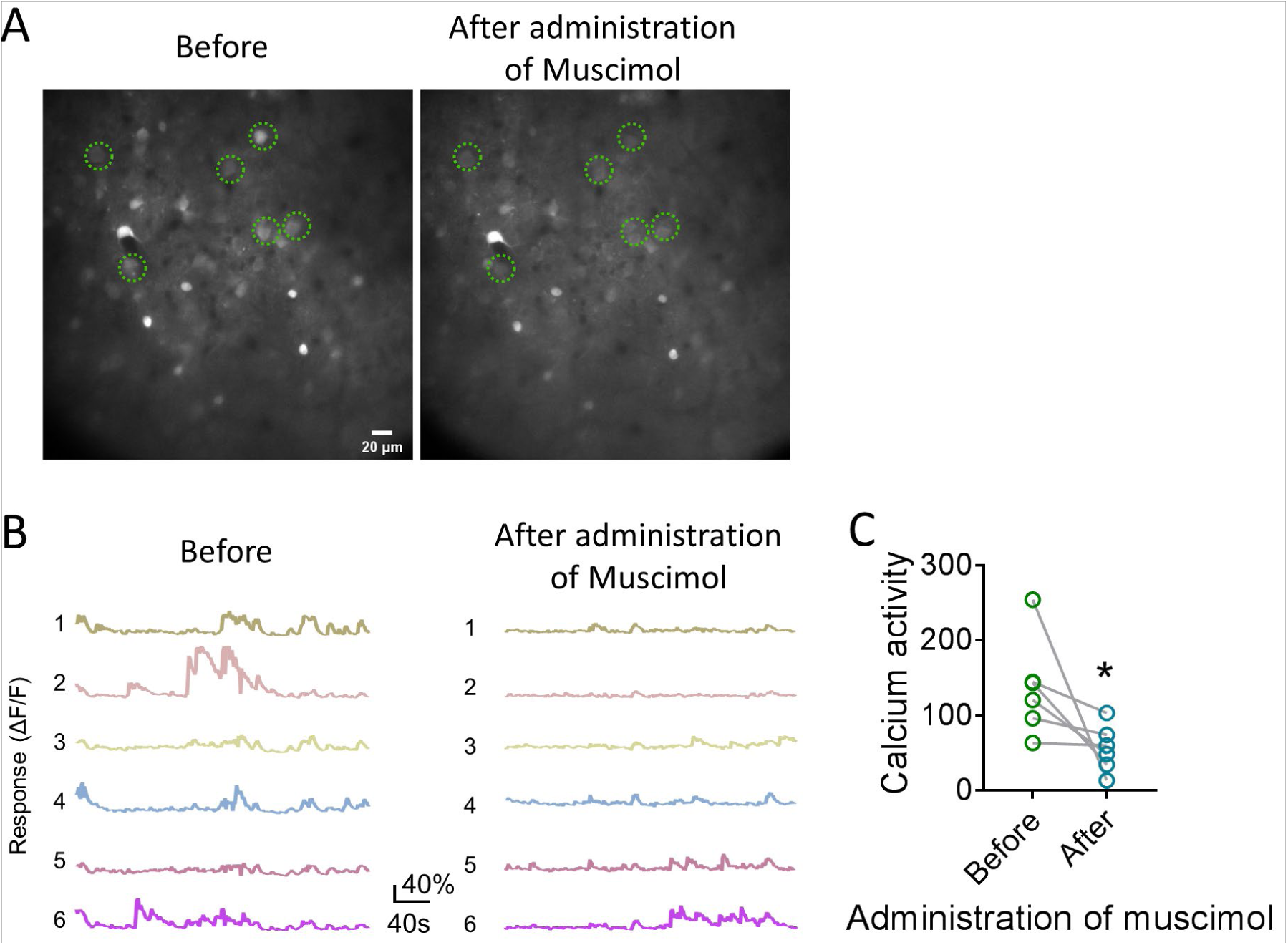
Neuronal network activity decreases after application of muscimol. (A) Two-photon time-lapse images of calcium activity (5 min intensity projection) in layer 2/3 excitatory neurons before and after intracerebral administration of muscimol (870 µM). (B) ΔF/F traces of six neurons before and after muscimol application. (C) Sum of neuronal ΔF/F (value>1.2) in 5 min (calcium activity) before and after application of muscimol. N=6 neurons, \**P*<0.05, paired t-test.

**Fig. S5.**
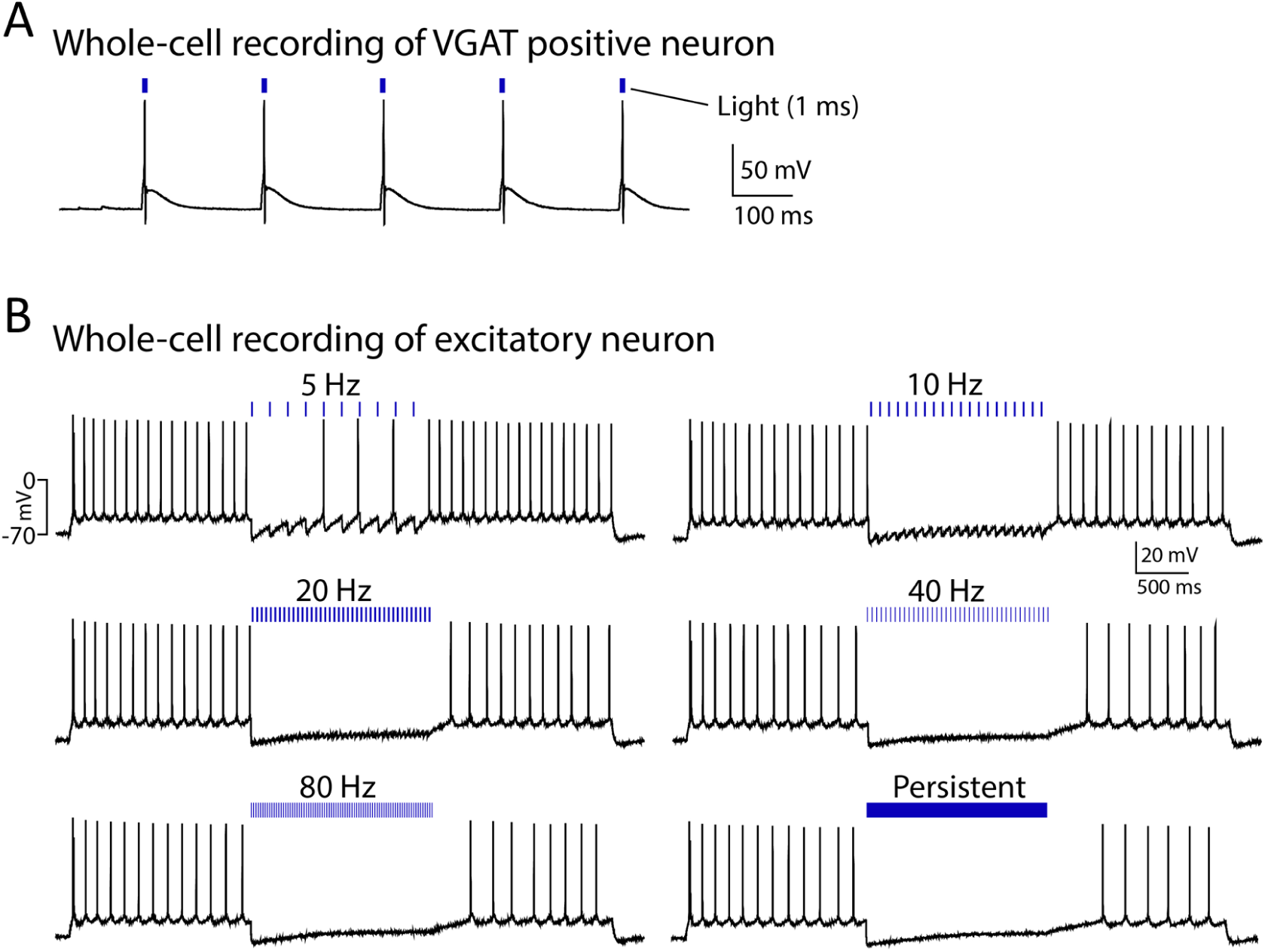
Verification of channelrhodopsin-2 expression in VGAT positive neurons and function in excitatory neurons. (A) Whole-cell recording in VGAT-positive neurons (inhibitory neurons) expressing ChR2 and 1 ms light induced action potential in VGAT-positive neurons. (B) Different frequencies of light stimulation inhibited firing in excitatory neurons.

**Fig. S6.**
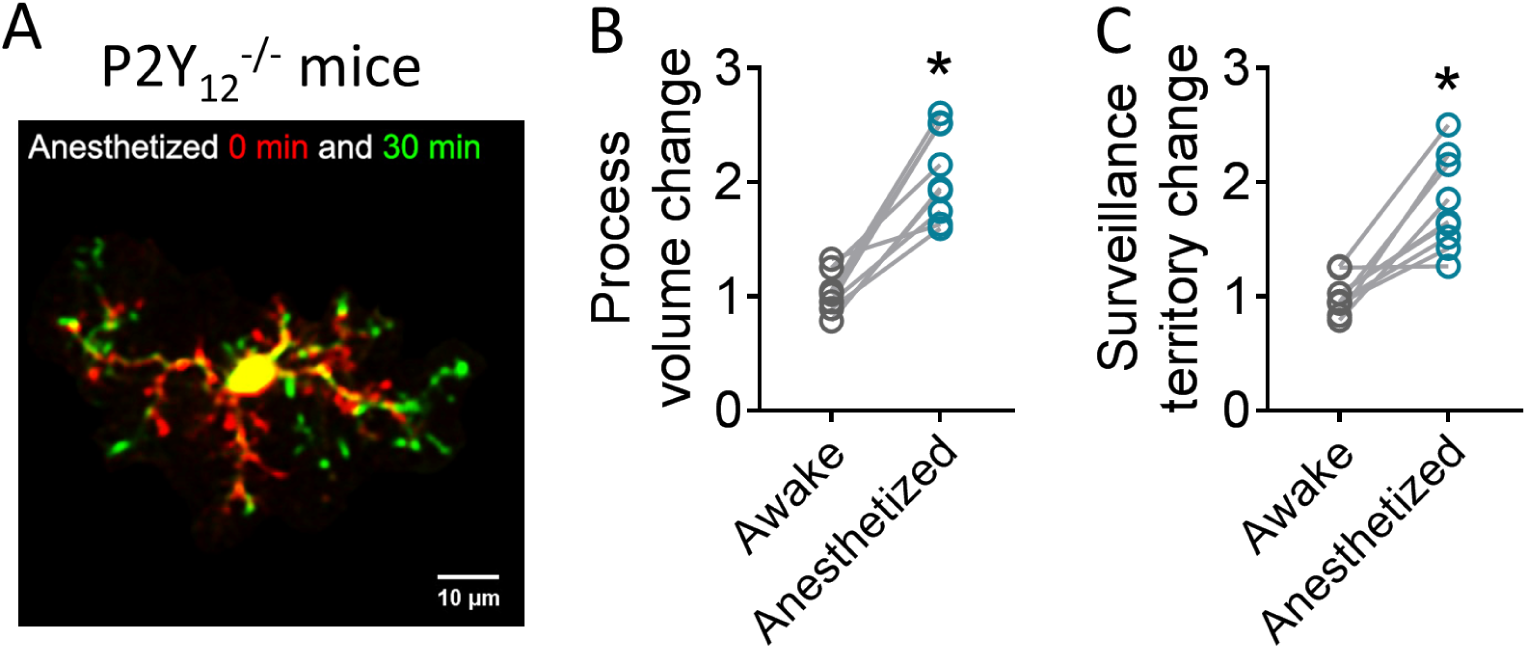
P2Y12 knockout does not affect the increase of microglial process surveillance by anesthesia. (A) Two-photon imaging of a microglia before (awake, red) and 30 min after (green) isoflurane anesthesia in P2Y_12_^-/-^ mice. (B and C) Microglial process dynamics were still increased in P2Y_12_^-/-^ mice after isoflurane anesthesia. All points represent 9 cells from 3 mice/group, \**P*<0.001, paired t-test.

**Fig. S7.**
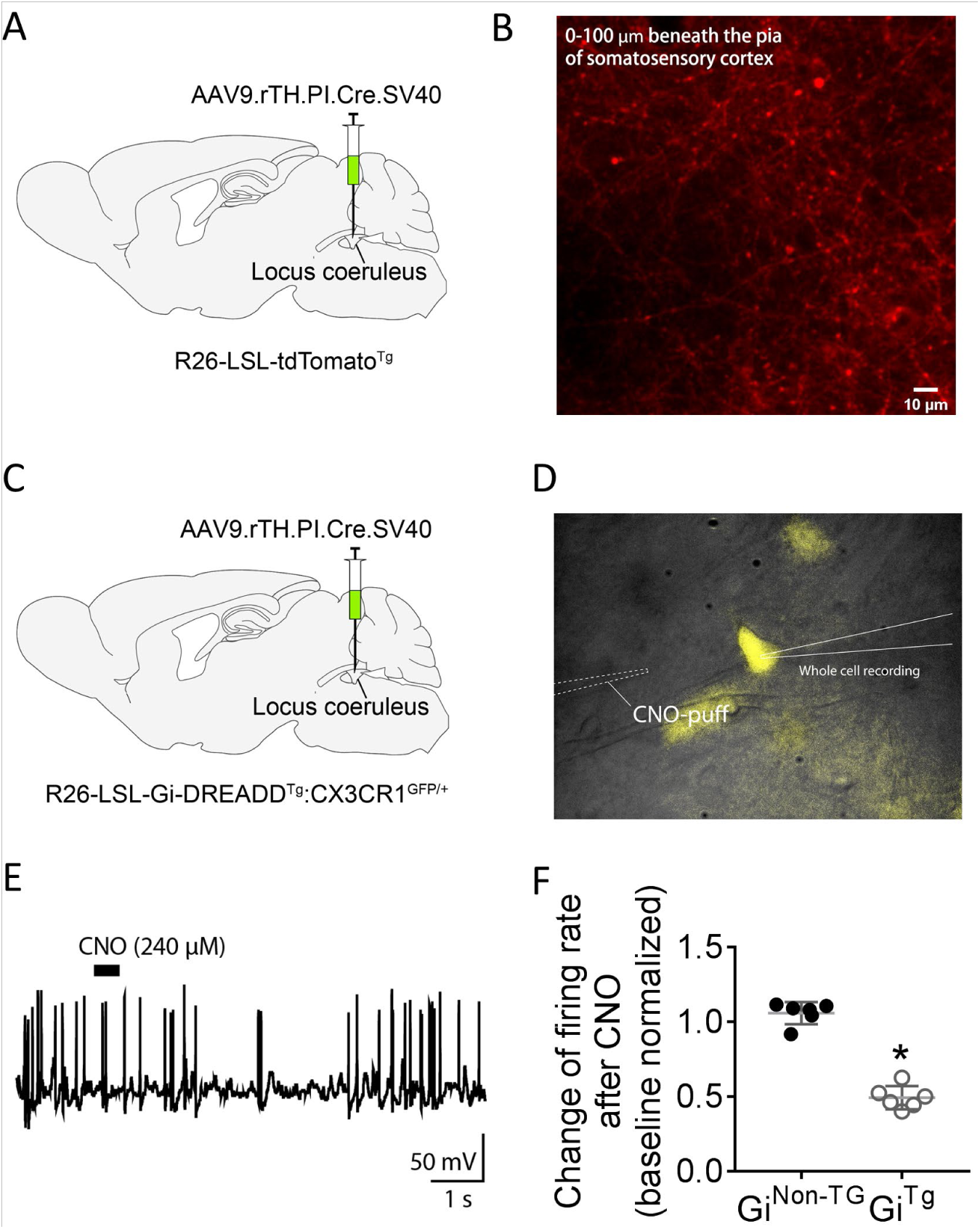
Adrenergic neurons of the locus coeruleus project to somatosensory cortex and functional expression of Gi-DREADD in the adrenergic neurons. (A and B) An adeno-associated virus, expressing Cre recombinase driven by tyrosine hydroxylase (rTH) promoter (AAV9.rTH.Cre), was injected in the locus coeruleus (LC) of a Cre reporter (tdTomato) mice (A). Two-photon imaging of tdTomato in somatosensory cortex of the reporter mice injected with the AAVs (Projection of 66 slices from 0-100 µm beneath the cortical surface) (B). (C and D) The AAV9.rTH.Cre was injected in the LC of mice allow Cre-inducible expression of Gi-DREADD and mCitrine yellow fluorescent protein (C). Whole-cell recording of mCitrine positive Gi-DREADD expressing adrenergic neurons in LC and glass pipette for CNO puffing (D). (E) Representative recordings of a Gi-DREADD expressing adrenergic neuron and the inhibition of neuronal firing by clozapine N-oxide (CNO). (F) Pool results showed CNO reduced firing in Gi-DREADD expressing neurons (Gi^TG^) but not in control neurons without Gi-DREADD expression (Gi^Non-TG^). \**P*<0.001, t-test.

**Fig. S8.**
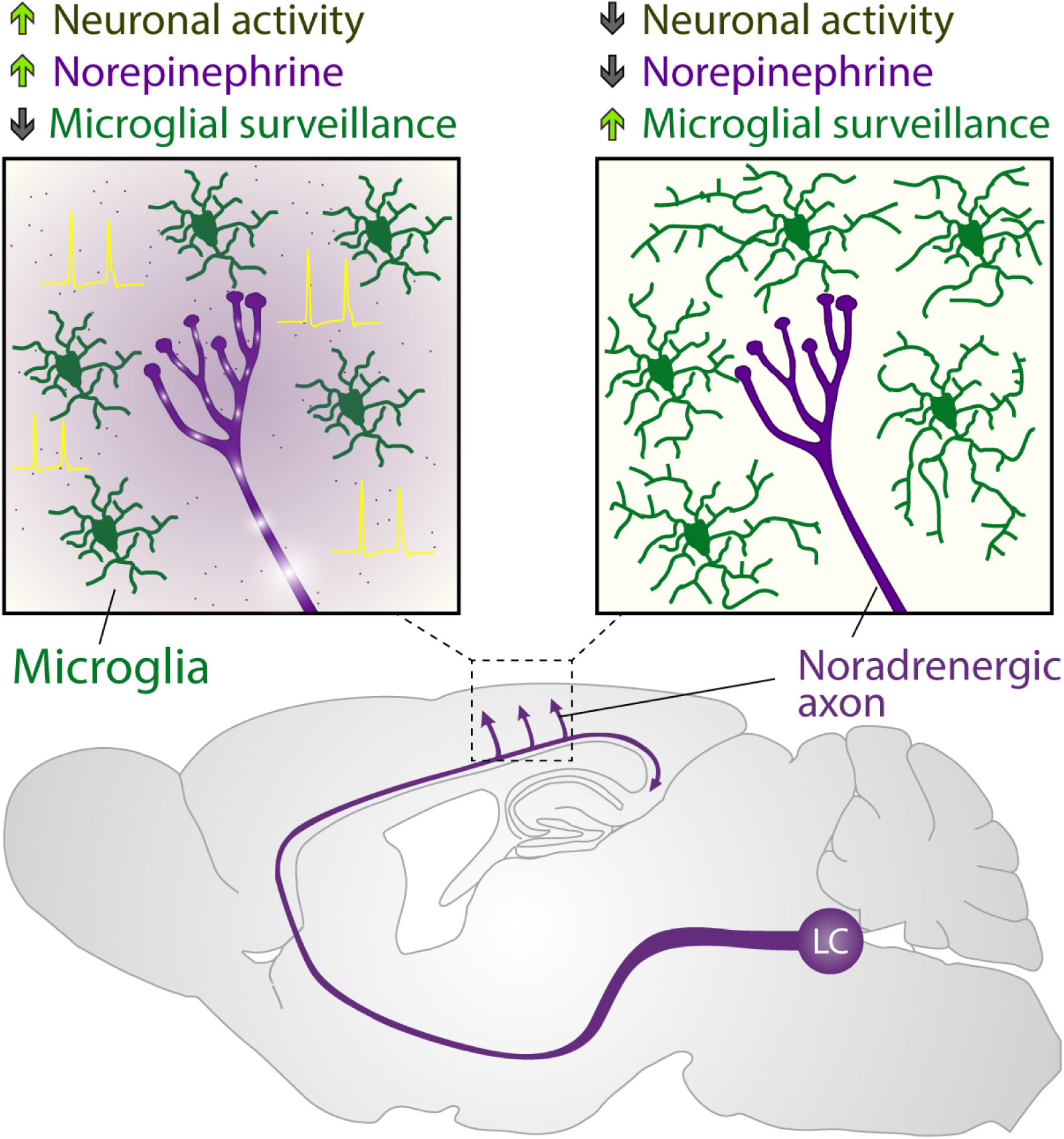
Neuronal network activity controls microglia process surveillance in awake mice via norepinephrine signaling. When neuronal activity is reduced by various manipulations, such as general anesthesia and sensory deprivation, microglial processes increase dynamics to survey more territory of the brain parenchyma due to reduced norepinephrine tone (purple).

**Movie S1. Microglial process surveillance increase after anesthesia.**

Time-lapse imaging of microglia at 50-64 µm beneath cortical surface in the somatosensory cortex before and after anesthesia.

**Movie S2. Neuronal network activity of the somatosensory cortex decrease after anesthesia.**

Calcium activity of excitatory neurons at 150 µm (left) or their dendrites at 50 µm (right) beneath the cortical surface of the somatosensory cortex before and after anesthesia.

**Movie S3. Neuronal network activity of the barrel cortex decreases after contralateral whisker trimming.**

Calcium activity of neuronal dendrites at 50 µm (right) beneath the cortical surface of the barrel cortex before and after whisker trimming.

**Movie S4. Microglia process surveillance increases after contralateral whisker trimming.**

Time-lapse imaging of microglia at 50-64 µm beneath cortical surface in the barrel cortex before and after anesthesia.

**Movie S5. Intracerebral administration of muscimol (870 µM) reduces neuronal network activity.**

Calcium activity of excitatory neurons at 150 µm beneath the cortical surface of the somatosensory cortex.

**Movie S6. Microglia process surveillance increases after intracerebral administration of muscimol (870 µM).**

Time-lapse imaging of microglia at 50-64 µm beneath cortical surface in the somatosensory cortex

**Movie S7. Microglia process surveillance increases after light stimulation (1ms, 10 Hz, 10 min) in mice that expressed channelrhodopsin 2 in VGAT-positive neurons (GABAergic neurons).**

Time-lapse imaging of microglia at 50-64 µm beneath cortical surface in the somatosensory cortex

**Movie S8. *In vivo* imaging of adrenergic neuronal projections from the locus coeruleus (LC)**

Imaging of axons from LC noradrenergic neurons which were expressing tdTomato, at 0-100 µm beneath cortical surface in the somatosensory cortex.

